# Regulation of adult neurogenesis and neuronal differentiation by Neural Cell Adhesion Molecule 2 (NCAM2)

**DOI:** 10.1101/2022.02.03.478938

**Authors:** Alba Ortega-Gascó, Antoni Parcerisas, Keiko Hino, Vicente Herranz-Pérez, Fausto Ulloa, Alba Elias-Tersa, Miquel Bosch, José Manuel García-Verdugo, Sergi Simó, Lluís Pujadas, Eduardo Soriano

## Abstract

Adult neurogenesis persists in mammals in the neurogenic zones where newborn neurons are incorporated into existing neuronal circuits. Relevant molecular elements of the neurogenic niches include the family of Cell Adhesion Molecules (CAM), which participate in signal transduction and regulate radial glial progenitor’s (RGPs) survival, division and differentiation. The Neural Cell Adhesion Molecule 2 (NCAM2) is expressed in brain development and in adult stages, and controls dendrite arborisation and synaptic formation and maintenance during development. Nevertheless, the role of NCAM2 in neurogenesis and lineage progression is not well understood. Here we analyse the functions of NCAM2 in the regulation of RGPs in adult neurogenesis in the dentate gyrus and during corticogenesis, by using different lentiviral-mediated genetic approaches to modulate its expression, both *in vivo* and *in vitro*. First, we characterized the expression of NCAM2 among the main actors of the neurogenic process revealing different levels of NCAM2 amid the progression of RGPs and the formation of juvenile neurons. Further, we show that overexpression of NCAM2 arrest infected cells in a RGP-like state, with characteristic morphological, immunocytochemical and electron microscopy features. In contrast, NCAM2 overexpression in embryonic cortical progenitors does not seems to alter cell fate, but causes transient migration deficits. These results reveal a differential role of NCAM2 in the regulation of adult and embryonic RGPs, and specifically, a significant implication of NCAM2 in the regulation and progression of RGPs during adult neurogenesis in the hippocampus.

## INTRODUCTION

In mammals, active neurogenesis is preserved during adulthood in specific niches (Altman and Das 1965) by remaining radial glia progenitors (RGPs) in the subventricular zone (SVZ) of the lateral ventricles and in the subgranular zone (SGZ) of the hippocampal dentate gyrus (DG) (Gonçalves et al. 2016; Gage 2019; Ghosh 2019; Kumar et al. 2019; Denoth-Lippuner and Jessberger 2021). Adult neurogenesis recapitulates the developmental processes including proliferation, neuronal fate specification, migration, differentiation, synaptogenesis, and functional integration into preexistent circuits. It has been shown that neurogenesis in the adult brain plays an important role in memory and learning processes (Zhao et al. 2008; Bergmann et al. 2015; Kumar et al. 2019).

RGPs are located in specialized microenvironments or neurogenic niches where they are subjected to multiple signaling pathways that control its maintenance, proliferation and lineage progression. Those extrinsic and intrinsic cues include cytokines, trophic and growth factors, neurotransmitters, epigenetic mechanisms as well as physiological and pathological variables (Yao et al., 2016; Zhang & Sheng et al., 2015; Zhang, 2018; Zhao et al., 2008). Cell adhesion molecules (CAMs) have also been revealed as essential components of these microenvironments. They not only sustain the cytoarquitechture of the niche but also provides a link between the extracellular and the intracellular domains of RGPs participating in signal transduction. Therefore, CAMs are important for self-renewal and proliferation of RGPs, and for neuronal differentiation and migration (Bian 2013; Morante-Redolat and Porlan 2019).

The mammalian neural cell adhesion molecule (NCAM) family is composed of two members, NCAM1 and NCAM2, sharing a similar structure of 5 immunoglobulin domains and 2 fibronectin type III domains, but presenting different expression patterns, post-transcriptional modifications, and molecular interactions (Pébusque et al. 1998; Makino and McLysaght 2010; Parcerisas, Ortega-gascó, et al. 2021). NCAM1 has been extensively studied and it has been described to play a role in neuronal migration, neurite development, synaptogenesis, and also in neurogenesis by regulating embryonic and adult neural stem cells (NSCs) (Kiselyov et al. 2003; Bonfanti 2006; Angata et al. 2007a; Boutin et al. 2009; Francavilla et al. 2009). NCAM2 has two different isoforms: NCAM2.1, with a cytoplasmatic domain, and NCAM2.2, which is GPI-anchored (Von Campenhausen et al. 1997; Alenius and Bohm 2003). In the central nervous system (CNS), the functions of NCAM2 have been mainly linked to the regulation of the formation and maintenance of axonal and dendritic biology compartments in the olfactory system (Alenius & Bohm, 2003; Kulahin & Walmod, 2010; Parcerisas, Ortega-Gascó, Pujadas, et al., 2021; Winther et al., 2012), and to the control of neural polarization, neurite outgrowth, dendrite development, and synapse formation and maintenance in the cortex and hippocampus through a complex panel of interactors (Leshchyns’Ka et al., 2015; Parcerisas et al., 2020; Sheng et al., 2015; Parcerisas, Ortega-gascó, et al., 2021). Interestingly, NCAM2 has been associated with different pathologies including Down syndrome, autism, and Alzheimer’s disease (JP et al., 2011; Leshchyns’Ka et al., 2015; Paoloni-Giacobino et al., 1997; Parr et al., 2006; Scholz et al., 2016; Winther et al., 2012). Regarding neurogenesis, Ncam2 has been detected in single-cell RNAseq studies that characterize the genetic profiles of qNSCs and their immediate progeny (Shin et al. 2015; Morizur et al. 2018). However, its role in RGP biology during neurogenesis remains unknown.

In the present study, we characterize the NCAM2 pattern of expression in the adult hippocampal neurogenic area and analyze the role of NCAM2 in the regulation of RGP biology during corticogenesis and in adulthood. To gain further insight into the importance of NCAM2 in the abovementioned processes, we used different biological and genetic tools including hippocampal viral injections, *in utero* electroporations and *in vitro* neurosphere cultures. Together, our results indicate that regulated NCAM2 expression levels are crucial for proper adult neurogenesis in addition to its relevant role during brain development. Moreover, we suggest that NCAM2 participates in the fine regulation of quiescency in hippocampal RGPs, a mechanism that could help explaining some pathologies that have been linked to NCAM2 such as Alzheimer’s disease which bear a complex phenotype including altered neurogenesis.

## MATERIALS AND METHODS

### Animals

All experimental procedures were carried out following the guidelines of the Committee for the Care of Research Animals of the University of Barcelona, in accordance with the directive of the Council of the European Community (2010/63 y 86/609/EEC) on animal experimentation. The experimental protocol was approved by the local University Committee (CEEA-UB, Comitè Ètic d’Experimentació Animal de la Universitat de Barcelona) and by the Catalan Government (Generalitat de Catalunya, Departament de Territori i Sostenibilitat).

### Antibodies and reagents

The following commercial primary antibodies were used for immunohistochemistry: Anti-ChFP (ab167453, Abcam, 1:300); Anti-DCX (A8L1U, Cell Signaling, 1:500) Anti-GFP (A11122, Invitrogen, 1:2000); Anti-GFAP (Z033401, DAKO, 1:2000); Anti-MAP2 (MA1406, Sigma, 1:2000); Anti-NCAM2 (AF778, R&D Systems, 1:750); Anti-Nestin (MAB353, Chemicon, 1:100), Anti-NeuN (MAB377, Merck, 1:1000); Anti-Sox2 (ab97959, Abcam, 1:500), Anti-Tbr2/EOMES (23345, Abcam, 1:100). Alexa Fluor fluorescent secondary antibodies were from Invitrogen. To counterstain nuclei, the tissue and cells were incubated in 2-(4-amidinophenyl)-1H -indole-6-carboxamidine (DAPI, D-6564, Sigma, 1:1000). Biotinylated-secondary antibodies were from Vector Labs; streptavidin-biotinylated/HRP complex and ECL were from GE Healthcare. The HRP-labeled secondary antibodies used for western blot were from DAKO. Diaminobenzidine reagent (DAB) and Eukitt mounting media were from Sigma-Aldrich. Mowiol was from Calbiochem.

### Plasmids

The plasmids ShNcam2, pCNcam2.1 and pCNcam2.2 used were described in Parcerisas et al, 2020. The cDNA of Ncam2.1 was amplified from the pCNcam2.1 with 5’-ACCATGAGCCTCCTCCTCTCC-3’ and 5’-CTGACCAAGGTGCTGAAACT-3’and cloned into pWPI (Plasmid #12254, Addgene) within PmeI site to obtain the pWPI-NCAM2.1. The cDNA of Ncam2.2 was amplified with 5’-ACCATGAGCCTCCTCCTCTCC-3’ and 5’-TCTCTGATCAGGGAGTACCA-3’ and cloned into pWPI (Plasmid #12254, Addgene) within PmeI site to obtain the pWPI-NCAM2.2.

### Production and intrahippocampal injection of retrovirus

The production and intrahippocampal injection of virus was performed as previously described (Parcerisas et al., 2020; Teixeira et al., 2012). Briefly, viral vectors were produced by transient transfection of HEK293T cells with calcium phosphate. Virus were concentrated by ultracentrifugation and resuspended in PBS.

For intrahippocampal injections, 8-week-old mice were anaesthetized with ketamine/xylazine mixture and placed on a heating blanket. They were positioned in a Kopf stereotaxic frame and a midline scalp incision was made. The scalp was reflected by hemostats to expose the skull, and bilateral burr holes were drilled. Viruses were then injected (1.5 μl of viral stock solution per site) into the left and right dentate gyrus over 20 min using a 5 µl Hamilton syringe. The micropipette was left in place for an additional 5 min. The coordinates used for the injections (in mm from Bregma and mm depth below the skull) were as follows: caudal 2.0, lateral 1.6, depth 2.2.

### Histological staining and electron microscopy

Animals were anaesthetized and perfused for 20 min with PBS 4% paraformaldehyde (PFA). The brains were then removed, post-fixed overnight with PBS 4% PFA, cryoprotected with PBS-30% sucrose and frozen. Coronal sections (30 μm) were obtained with a cryostat and immunohistofluorescence or immunohistochemistry were performed on free-floating sections. Samples were blocked with PBS containing 10% normal horse serum (NHS) and 0.2% gelatin; and incubated at 4^°^C overnight with PBS-5% NHS primary antibodies. For immunohistofluorescence, sequential incubation was carried out using a secondary antibody (Alexa Fluor, Invitrogen), and the sections were mounted with Mowiol (Calbiochem). The images were acquired with confocal microscopy (Spectral Confocal SP2 Microscope, Leica; Spectral Confocal SP8, Leica and Carl Zeiss LSM880, Zeiss). For immunohistochemistry, sequential incubation was carried out using biotinylated secondary antibodies (2 h at room temperature) and streptavidin-HRP (1:400; 2 h at room temperature) was performed in PBS-5% NGS; bound antibodies were visualized by reaction using DAB and H_2_O_2_ as peroxidase substrates; the sections were dehydrated and mounted (Eukitt). Images were acquired with AF6000 microscope (Leica) and Olympus Bx61 microscope (Olympus).

For electron microscopy, sections were cryoprotected in 25% saccharose and freeze-thawed (3×) in methylbutane. The sections were then washed in 0.1 M phosphate buffer (PB; pH 7.4), blocked in 0.3% bovine serum albumin-C (BSA), and incubated with a primary chicken anti-GFP antibody (1:200; Aves Labs, Tigard, OR, USA) for 72 h at 4^**°**^C. The sections were washed in phosphate buffer (PB), blocked in 0.5% BSA and 0.1% cold-water fish-skin gelatin (Electron Microscopy Sciences, Hatfield, PA, USA) for 1 h, and subsequently incubated with a colloidal gold-conjugated secondary antibody (1:50; goat anti-chicken IgG gold UltraSmall, Electron Microscopy Sciences) for 24 h at room temperature. The sections were then washed in PB and 2% sodium acetate. Silver enhancement (Aurion R-gent Silver enhancer kit, Electron Microscopy Sciences) was performed following the manufacturer’s directions, and the sections were washed again in 2% sodium acetate. To stabilize the silver particles, the samples were immersed in 0.05% gold chloride (Sigma) for 10 min at 4^**°**^C, washed in sodium thiosulfate, washed in PB, and then postfixed in 2% glutaraldehyde for 30 min. The sections were incubated in 1% osmium tetroxide and 7% glucose and then washed in deionized water. Subsequently, sections were partially dehydrated in 70% ethanol and incubated in 2% uranyl acetate in 70% ethanol in the dark for 2.5 h at 4^**°**^C. Brain slices were further dehydrated in ethanol followed by propylene oxide and infiltrated overnight in Durcupan ACM epoxy resin (Fluka, Sigma-Aldrich, St. Louis, USA). The following day, fresh resin was added, and the samples were cured for 72 h at 70^**°**^C. Following resin hardening, 1.5-µm semi-thin sections were selected under light microscopy based on their immunolabeling and detached from glass-slides by repeated freezing and thawing in liquid N_2_. Ultra-thin sections were obtained at 60–70 nm from selected semi-thin sections. Photomicrographs were obtained using a FEI Tecnai G^2^ Spirit (FEI Europe, Eindhoven, Netherlands) using a digital camera Morada (Olympus Soft Image Solutions GmbH, Münster, Germany).

### *In utero* electroporation

*In utero* microinjection and electroporation were performed at E14.5 as described (Simó et al. 2010; Parcerisas et al. 2020b), using timed pregnant CD-1 mice (Charles River Laboratories). Briefly, DNA solutions were mixed in 10 mM Tris (pH 8.0) with 0.01% Fast Green. Needles for injection were pulled from Wiretrol II glass capillaries (Drummond Scientific) and calibrated for 1 µl injections. Forceps-type electrodes (Nepagene) with 5-mm pads were used for electroporation (five 50-msec pulses of 45 V at E14.5). Brains were collected at E19.5/P0 or P5, dissected, and successful electroporations identified by epifluorescence microscopy. Positive brains were fixed in 4% formalin in 0.1 M phosphate buffer saline (PBS) and cryoprotected in 30% sucrose/PBS overnight at 4^°^C. Brains were frozen in O.C.T compound before fourteen-micrometer-thick brain cross-sections were obtained with cryostat and placed on slides. Sections were antigen-retrieved by immersion of the slides in 0.01 M sodium citrate buffer, pH 6.0 at 95^°^C for 20 min. Sections were blocked for 2 h with 10% normal goat serum, 10 mM glycine, and 0.3% Triton X-100 in PBS at room temperature. Primary antibodies (anti-GFP and anti-ChFP) were incubated overnight at 4^°^C. Slides were washed four times for 10 min in 0.1% Triton X-100/PBS. Secondary antibodies were added for 2 h at room temperature and the slides were washed as before and coverslipped with Prolong Gold anti-fade reagent (Molecular Probes). Most images were obtained with epifluorescent illumination and a 10× objective (Leica 760 or AF6000). Positions of GFP-or ChFP-positive neurons were recorded from several sections per embryo. Data were collected from the lateral part of the anterior neocortex. For a BIN10 quantification, the cortex was divided into ‘BINs’ as follows: the distance from the pial surface to the bottom of the SVZ was measured and divided into 10 equal-sized BINs. The percentage of GFP-or ChFP-labeled neurons in each BIN for each embryo was then calculated. Graphs plot the mean and standard error of % neurons in each BIN for the N embryos in a group.

### Neurospheres culture

Neurospheres cultures were derived from 7-8 postnatal day (P7-P8) mice following the modified protocol described by Walker & Kempermann, 2014. Briefly, the SVZ of the lateral ventricles and the SGZ of the hippocampus were dissected in PBS. After trypsin (GIBCO) and DNAse (Roche diagnostic) treatments, the tissue was dissociated with gentle sweeping. Cells were counted and plated in non-adherent 24 well plates in Neurobasal medium containing 2% B27 supplement (GIBCO), penicillin/streptomycin (Life technologies) and Glutamax (Life technologies), 20 ng/ml EGF, 20 ng/ml bFGF and 2 μg/ml heparin. Cells were incubated at 37^**°**^C with 5% CO_2_ and subcultured every 2-3 days.

For the growth analysis, neurospheres from the SGZ were dissociated with trypsin and infected at passage 2 with viruses (pWPI, pWPI-NCAM2.1, pWPI-NCAM2.2, ShNcam2 or ShCnt). GFP positive cells were selected by flow cytometry (BD FACSAria Fusion), plated in non-adherent 24-well plates and analyze during 5 consecutive days. High content image acquisition was performed with an Automated Wide-field Olympus IX81 Microscope (Olympus Life Science Europe, Waltham, MA) and a 4x UPlan FL N objective. ScanR Acquisition software version 2.3.0.5 was used to automatically record adjacent fields of view taking 20 (5 × 4) z-stacks (8 slices with a z-step of 200 nm) per well, with 10% of overlap to enable automatic image stitching. Neurosphere size was quantified by means of 3 different Fiji macros. In brief, tailor-made macros were used to project each z-stack, to stitch these projections and to quantify the size of each neurosphere.

## Image analysis

All images were processed and quantified using the ImageJ software (NIH).

## Statistical analysis

Statistical analysis was carried out using the Prism 8 software. Significance between two experimental groups was analysed using the unpaired Student’s *t*-test. Differences between groups in distribution of cells in corticogenesis were assessed by two-way ANOVA followed by Bonferroni’s comparison *post hoc* test. To determine differences between more than two groups in the adult neurogenesis characterization experiments, one way ANOVA was used. Post-hoc comparisons were performed by Tukey’s test and significance level was set at P>0.05: *P<0.05, **P<0.01, and ***P<0.001. To determine differences between two groups, Student’s t-test and significance level was set at P>0.05: *P<0.05, **P<0.01, and ***P<0.001. Statistical values are presented as mean ± standard error of the mean (SEM).

## RESULTS

### Differential expression pattern of NCAM2 in the dentate gyrus

Since cell adhesion molecules are important structural elements of the neurogenic niches we first characterize the expression of NCAM2 in the different populations of cells at the DG of P45 mice by immunofluorescence. Dentate RGPs undergo several morphological and electrophysiological changes while expressing different markers through the neurogenic process to finally give rise to mature neurons. To identify type I progenitors we used the GFAP/Sox2 or Nestin markers while Tbr2 was selected to mark type II proliferative progenitors (Kempermann et al., 2015). In addition, we detected neuroblasts or immature neurons with antibodies against DCX; and mature neurons labelling NeuN. As NCAM2 is a membrane protein, the general pattern of NCAM2 staining show clear staining in the delineating cells bodies and the dendrites of neurons. Confocal microscopy analysis reveal strong NCAM2 signal in GFAP/Sox2 or Nestin positive cells with the typical morphology of type I progenitors (i.e: triangular cell body located in the SGZ and a unique dendrite extended into the molecular layer) (**Fig. 1A-B**). Contrariwise, images suggest that NCAM2 expression in Tbr2 positive cells is low, although it is difficult to determine the expression of NCAM2 and Tbr2 in the same cells due to the localization of both proteins (**Fig. 2C**). Among the DCX positive cells population, we found different phenotypes with differences in NCAM2 staining. While some DCX positive cells display faint or undetected NCAM2 staining, other cells present higher levels of the protein (**Supplementary Fig. 1A**). Lastly, mature granule cells that express NeuN also present NCAM2 labelling, as expected (**Supplementary Fig. 1B**).

**Figure 1.**
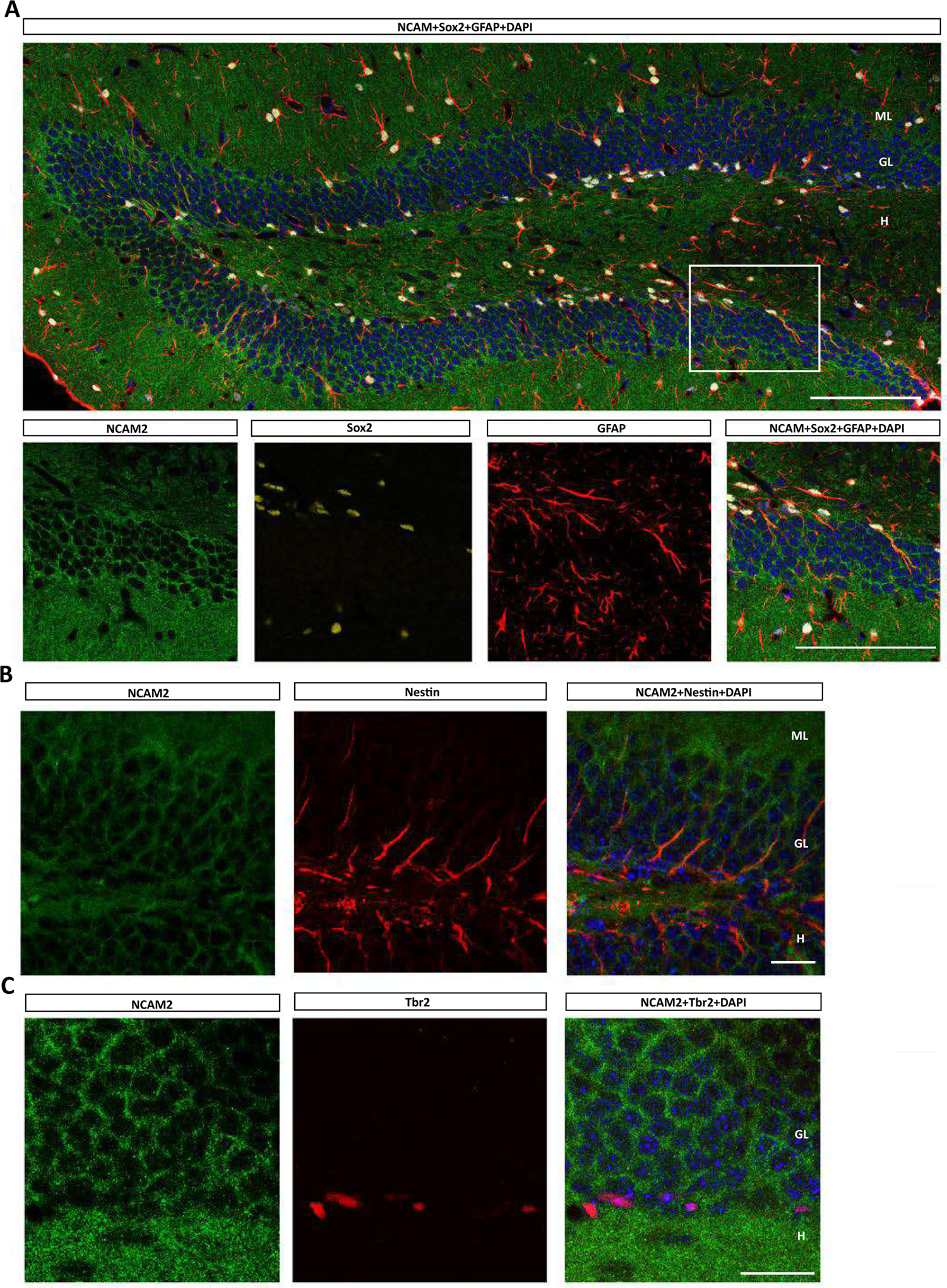
Expression pattern of NCAM2 in the hippocampus progenitor cells. **A)** Immunohistochemical characteritzation of NCAM2 expression in GFAP/Sox2 progenitor cells in P45 mice hippocampus. Arrowheads label NCAM2/GFAP/Sox2 positive cells. **B) N**CAM2 expression in Nestin positive cells in the subgranular zone of P45 mice. Arrowheads label NCAM2/Nestin cells. **C)** Double immunostaining of NCAM2 and Tbr2 at P45. Arrowheads label Tbr2 positive cells that present low NCAM2 signal. ML: molecular layer; GL: granule layer; H: hilus. Scale bar: A) 50 μm, B,C) 20 μm.

**Figure 2.**
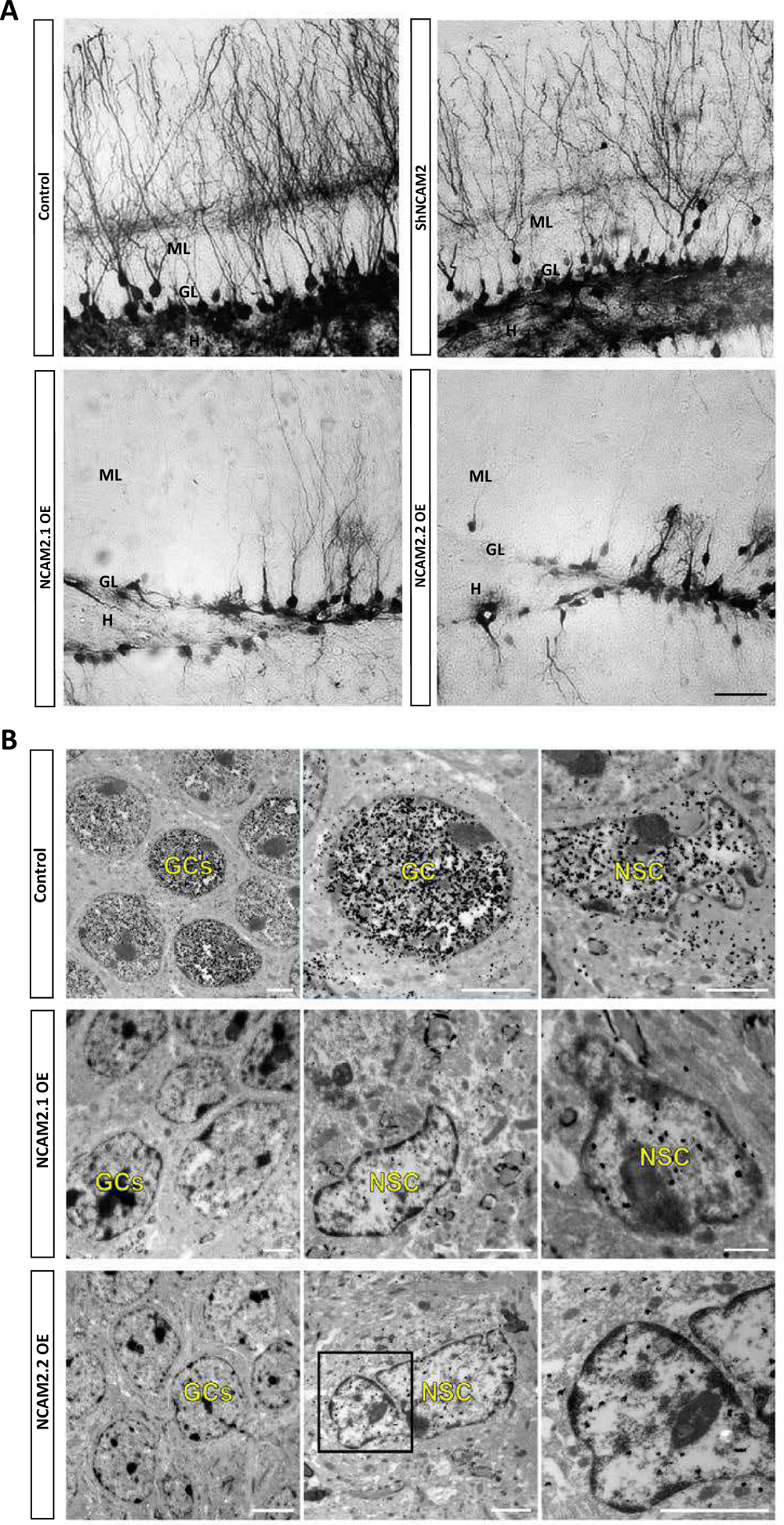
NCAM2 overexpression modulates adult neurogenesis in the hippocampus. **A)** Representative images of GFP positive cells from the dentate gyrus of mice injected with control, ShNCAM2, or NCAM2 overexpressing viruses (NCAM2.1 and NCAM2.2) at 4 weeks after injection. Control and ShNCAM2 positive cells show a granule cell morphology while RGP-like phenotype was observed in many cells infected with NCAM2.1 or NCAM2.2 overexpressing viruses. Scale bar: 50 µm. **B)** GFP immunogold electron microscopy images of animals infected with control, NCAM2.1 OE or NCAM2.2 OE viruses and sacrificed 4 weeks post-surgery. Control images show densely GFP-labelled granule cells and RGPs. In NCAM2.1 and NCAM2.2 OE mice, the number of labelled granule cells is dramatically decreased. Nevertheless, RGPs located in the SGZ still appear labelled with GFP. GC: granule cell; RGP: radial glia progenitor. Scale bar: 2 µm.

Therefore, the characterization of the expression pattern of NCAM2 in the dentate gyrus of the hippocampus suggests a differential expression of the protein in the main actors of the neurogenic process: while both RGPs and mature neurons express appreciable NCAM2 staining the intermediate type II-III progenitors may have a minimum in NCAM2 expression.

### NCAM2 modulates adult neurogenesis in the hippocampus

With the purpose to study the potential role of NCAM2 in adult neurogenesis, we modulate the expression of the NCAM2 protein in the hippocampal neurogenic region. We stereotaxically injected transduced the DG of 8 week-old mice with NCAM2.1/NCAM2.2-overexpressing or ShNCAM2-silencing lentiviruses, which bear preferential infectivity on progenitor cells or neuroblasts. We analyzed the transduced DGs 4 weeks after surgery. Mice injected with control viruses exhibited the characteristic morphology of dentate granule cells (i.e. round soma in the granule cell layer, and apical dendrites ramifying in the molecular layer and covered by dendritic spines) (**Fig.2A**, first panel). Similar results were found in mice injected with ShNCAM2 viruses, indicating that the downregulation of NCAM2 does not alter the formation, survival, or maturation of new adult-born neurons in the DG. (**Fig. 2A**, second panel). Conversely, we found that many cells infected with NCAM2.1 and NCAM2.2 overexpressing viruses did not exhibit the typical morphology of maturing granule cells but a RGP-like phenotype (i.e. triangular cell bodies located in the inner GL, with a unique, short radial process spanning the GL and ramifying profusely in the inner molecular layer) (**Fig. 2A**, third and fourth panel). Some infected cells, however, resembled type II progenitors or neuroblasts (i.e. irregular soma with short processes oriented tangentially or rounded soma with a short apical dendrite oriented towards the molecular layer) or immature granule cells. Enrichment in RGP-like phenotype apparently was more prominent upon NCAM2.2-overexpression.

To further characterize the phenotype of NCAM2 overexpressing cells, we performed fine structure analysis of GFP-labelled cells, identified by GFP-immunogold electron microscopy (**Fig. 2B**). Confirming our optical microscopy results, most control infected cells at the injection site corresponded to dentate granule cells which were closely apposed in the granule layer (GL). These cells showed a typical round-shaped soma, most of it occupied by the nucleus, which displayed chromatin aggregates. The cytoplasm was comprised by a thin space with a few long cisternae of endoplasmic reticulum and abundant free ribosomes. Nevertheless, we also observed GFP-positive cells in the SGZ. Among them, we identified RGPs and type II cells or neuroblasts. As previously described (Seri et al. 2004), RGPs were recognized as cells with a large cell body with a major radial process that penetrates the granular layer extending thin lateral processes between granule neurons. They present a round or triangular nucleus, electron lucent cytoplasm, irregular contour and intermediate filaments in their cytoplasm. On the other hand, type II cell (or neuroblast) features include a smooth contour, dark scant cytoplasm, abundant polyribosomes and a less developed endoplasmic reticulum than granule cells. Interestingly, NCAM2.1/NCAM2.2-overexpressing GFP-positive cells were mainly detected in the SGZ and, according to their fine structure, could be identified RGPs (**Fig. 2B**).

### Cell autonomous overexpression of NCAM2 retains adult-born DG cells in a RGP-like phenotype

To better understand the events triggered by the expression of NCAM2 isoforms, animals injected with control and NCAM2 overexpressing viruses were sacrificed at different time points including earlier stages (3 days, 1 week, 2 weeks and 4 weeks) (**Fig. 3A**). As a starting point, animals were sacrificed 3 days after injection. Although the infection is not strictly restricted to progenitor cells, as expected, the majority of the infected cells exhibit a RGPs morphology 3 days post-injection in all the experimental conditions (Consiglio et al. 2004; Jandial et al. 2008) (**Fig. 3B**). Focusing on posterior time points, most cells infected with control vectors at 1 week post-injection showed a morphology typical of immature granule cells that appeared progressively more mature at 2 and 4 weeks post-injection. In contrast, the shapes of NCAM2.1- and NCAM2.2-overexpressing cells remained constant overtime, with most of the labeled cells exhibiting an RGP-like cell morphology, while others exhibited intermediate progenitors-or neuroblast-like phenotypes (**Fig. 3B)**.

**Figure 3.**
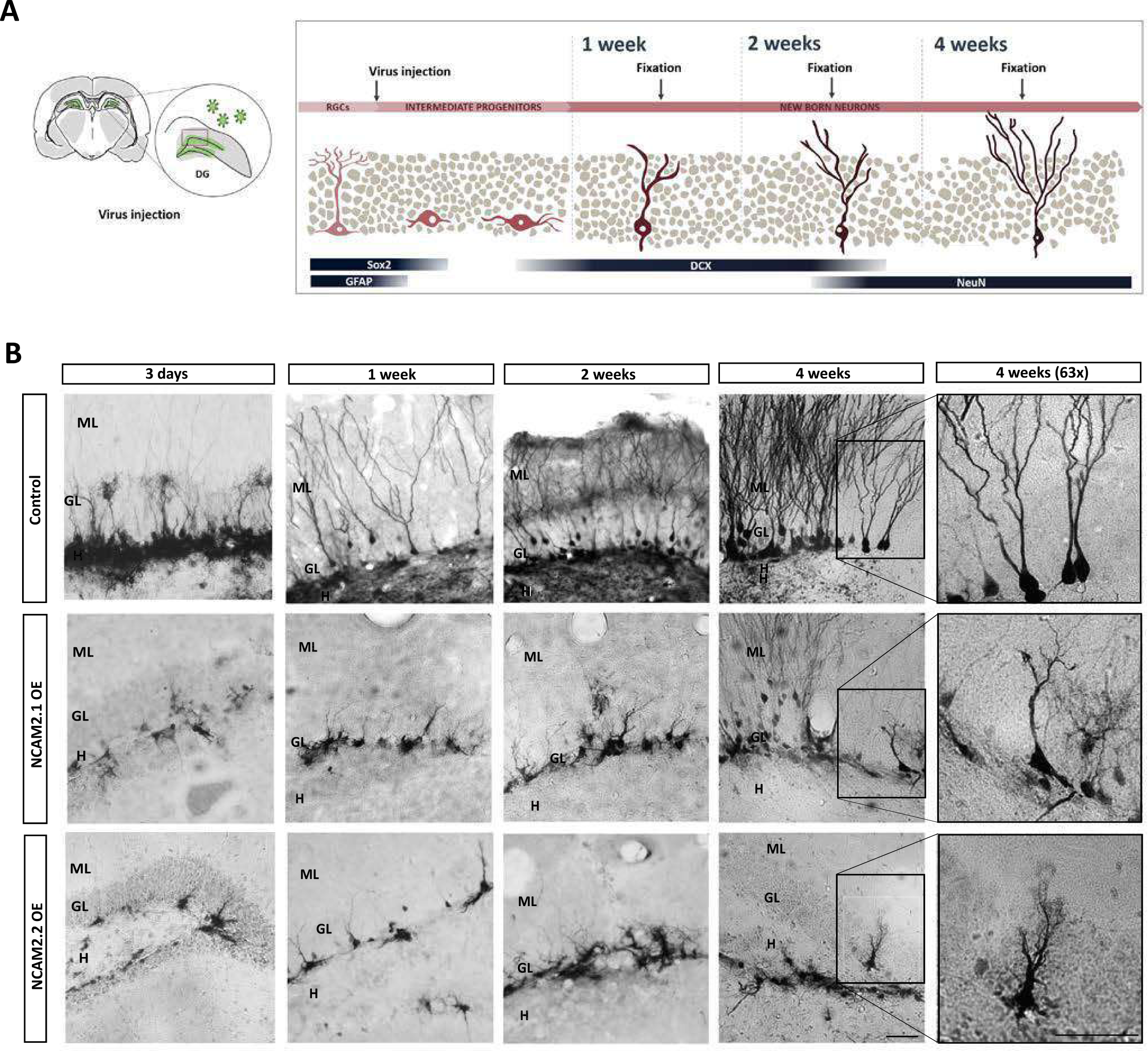

The phenotype of cells infected with the viral vectors was additionally characterized evaluating the expression of specific cell markers at the different time points analyzed. The triple immunostaining of GFP/Sox2/GFAP was used to determine the proportion of RGPs within the pool of infected cells (**Fig. 4A-B**). At 3 days after injection most of control-infected cells were Sox2/GFAP double positive (**Fig. 4C**). Similarly, also NCAM2.1- and NCAM2.2-infected cells were mostly positive for both markers at 3 days post infection (**Fig. 4C**). Analyzing the evolution of GFP-/Sox2-/GFAP-positive progenitors in the control conditions we observed a significant and progressive decline in the number of progenitors over time (**Fig. 4C-D**). In contrast, in the animals infected with NCAM2.1 or NCAM2.2 overexpressing viruses, we noticed a much less marked decrease in the proportion of those progenitors along the time-course, thus suggesting an arrest of the cells in the progenitor stage (**Fig. 4C-D**). We confirmed that most NCAM2.1- and NCAM2.2-overexpressing cells morphologically characterized as neuronal progenitor cells expressed the neuronal progenitor markers GFAP and Sox2 at 1 week after injection (**Fig. 4A,C)**. Additionally, quantification of the percentage of GFP/Sox2/GFAP revealed a maintenance of high proportion of GFAP/Sox2 positive cells in both NCAM2.1 and NCAM2.2 overexpressing conditions also at 2 weeks and 4 weeks, being more pronounced in the case of NCAM2.2 isoform (**Fig. 4C**).

**Figure 4.**
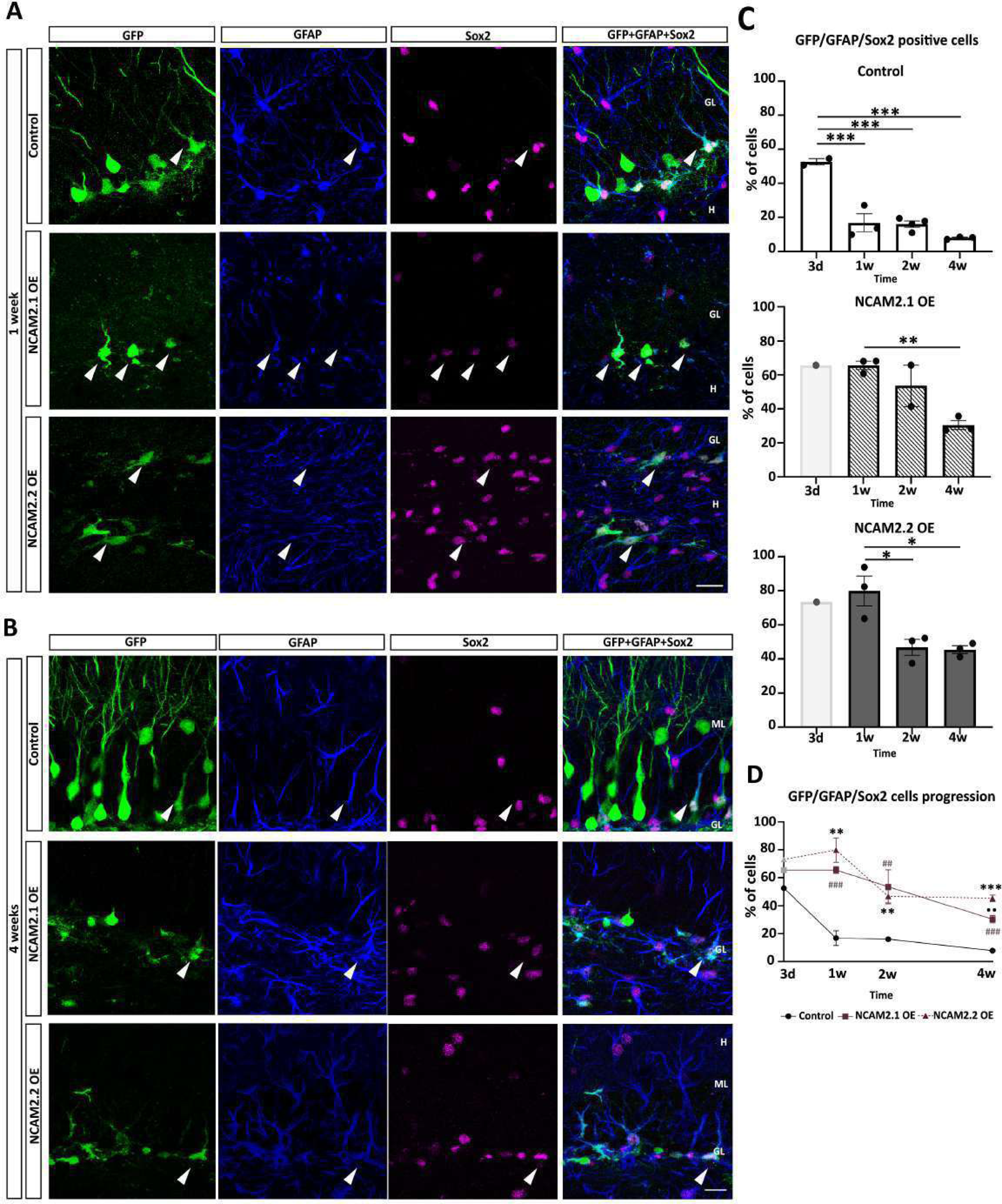
Immunohistochemical characterization of NCAM2 overexpressing progenitor cells. **A)** Immunostaining of GFP positive cells with GFAP and Sox2 RGPs markers from animals sacrificed 1 week post-injection. **B)** Immunostaining of GFP positive cells with GFAP and Sox2 RGPs markers from animals sacrificed 4 weeks post-injection. **C-D)** Time course quantification of the GFP/Sox2/GFAP positive cells in mice injected with control, NCAM2.1 OE or NCAM2.2 OE viruses at 3 days, 1 week, 2 weeks and 4 weeks post-injection. N=2-3 animals, 5-10 slices per animal. Data are presented as mean ± SEM; dots represent average values for individual animals (5-10 slices per animal, 20-50 cells per animal); N=2-3 animals per group at 3 days (control) 1, 2 and 4 weeks post-injection; ANOVA, Tukey’s comparison *post-hoc* test; * P<0.05, ** P<0.01, *** P<0.001, **** P<0.0001. In light gray bars, representation of NCAM2.1 and NCAM2.2 groups at 3 days post-injection (N=1 animals per group, qualitative study excluded form statistical analysis). In Fig. 4D, gray * differences between Control and NCAM2.1; black * differences between Control and NCAM2.2; • differences between NCAM2.1 and NCAM2.2. Arrowheads label GFP/Sox2/GFAP positive GFP-cells. ML: molecular layer; GL: granule layer; H: hilus. Scale bar: A, B) 20 µm.

The impact of NCAM2 overexpression in the process of neurogenesis was complemented quantifying the number of DCX positive cells at 2 and 4 weeks after viral transduction (**Fig. 5A**). According to the expected evolution of the neurogenic events, we observed a high percentage of DCX positive cells at 2 weeks after injection followed by a decrease at 4 weeks (**Fig. 5C**). The above-mentioned decline is not detected when NCAM2.1 or NCAM2.2 are overexpressed and we found a persisting number of DCX positive cells from 2 to 4 weeks after transduction. Finally, the number of NeuN mature neurons 4 weeks post-injection was also analyzed (**Fig. 5B**). In agreement with the previous data, we found a trend to show reduced percentages of NeuN positive neurons in the overexpression conditions at 4 weeks post-induction, reaching statistically significance for the NCAM2.2 isoform compared to controls (**Fig. 5D**).

**Figure 5.**
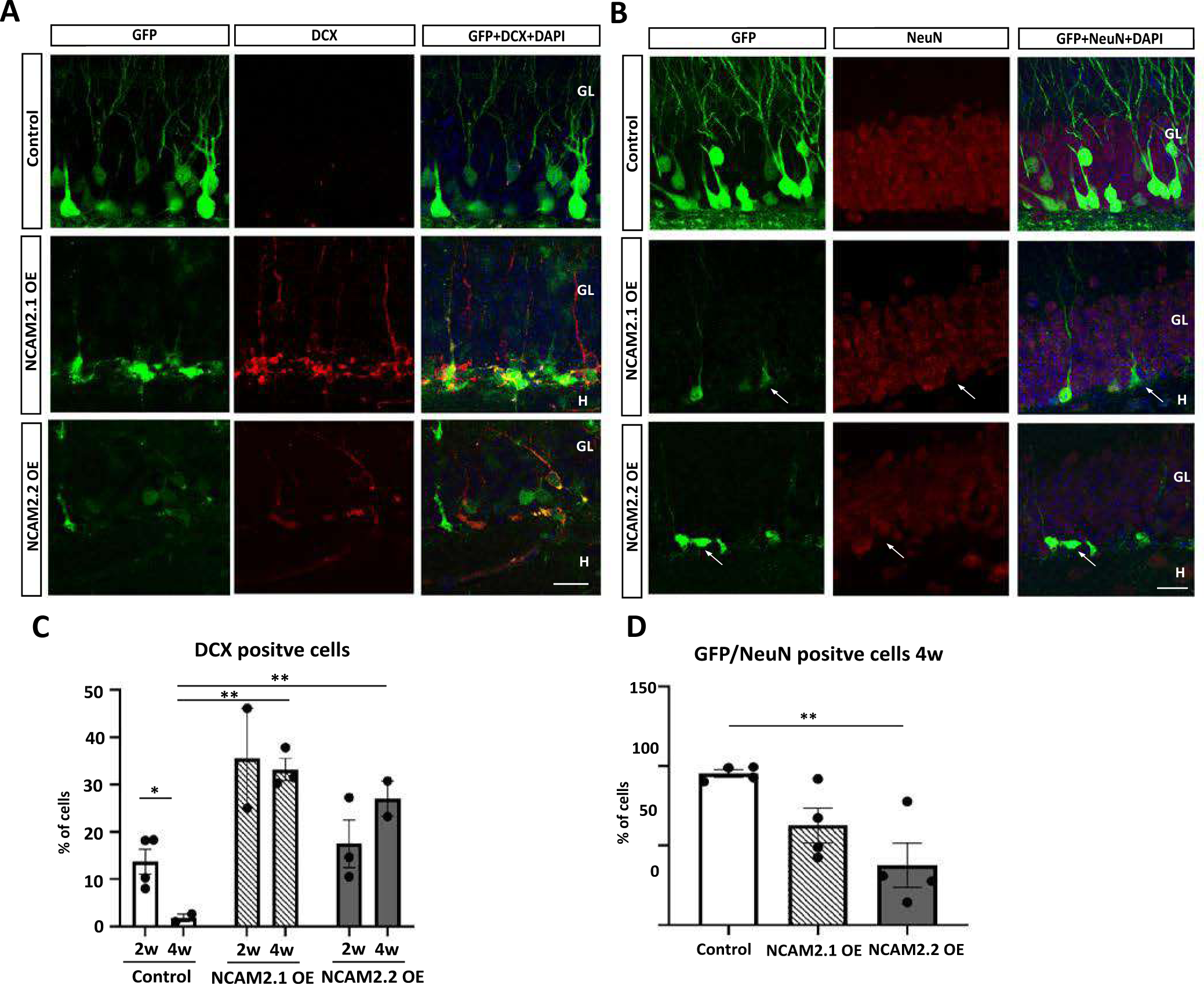
Immunohistochemical characterization of NCAM2 overexpressing neurons. **A)** Immunostaining of GFP positive cells with DCX as a markers for neuroblasts (type III progenitors) and immature neurons from animals sacrificed 4 week post-injection. **B)** Immunostaining of GFP positive cells with NeuN as a markers for mature neurons from animals sacrificed 4 week post-injection. **C)** Quantifications of GFP/DCX positive cells in mice injected with control, NCAM2.1 OE or NCAM2.2 OE viruses at 2 and 4 weeks post-injection. N=2-4 animals per group, 5-6 slices (>50 cells per animal in the controls; 15-30 cells per animal in the NCAM2 OE conditions). **D)** Quantification of GFP/NeuN positive cells in animals injected with control, NCAM2.1 OE or NCAM2.2 OE 4 weeks after transduction. N=4 animals per group, 5 slices per animal (>50 cells per animal). Data are presented as mean ± SEM; differences between experimental groups ANOVA, Tukey’s comparison *post-hoc* test; ** P<0.01; differences between time points Student’s t-test; * P<0.05. Arrowheads label DCX or NeuN positive GFP-cells; arrows label NeuN negative GFP-cells. ML: molecular layer; GL: granule layer; H: hilus. Scale bar: A, B) 20 µm.

This time-course analysis suggests that the observations at 4 weeks post-injection time on overexpression of NCAM2 are not attributable to a de-differentiation of immature neurons, but rather to a temporarily arrest of the RGP-like phenotype in the SGZ that leads to a delay in the formation of new neurons.

### NCAM2 overexpression do not arrest embrionary RGPs

Since our results point out to an important role of NCAM2 in the regulation of adult RGPs and NCAM1 is involved both in adult and embryonic neurogenesis (Angata et al. 2007b; Boutin et al. 2009), we next sough to study the potential role of NCAM2 in RGPs during embryonic stages. We performed *in utero* electroporation experiments using isoform-specific overexpressing vectors (i.e. NCAM2.1 and NCAM2.2). Embryos were electroporated at E15 (using GFP or ChFP as reporter genes) and brains were analysed at P0 and P5, by counting the distribution of electroporated neurons across cortical layers. Interestingly, we found a moderate non-significant of cells located in the neurogenic areas, VZ and the intermediate zone (IZ), in cortices electroporated with NCAM2 isoforms (**Fig. 6A**). However, we observed alterations in the migration of neurons when modulating NCAM2 expression. Our previous study (Parcerisas et al., 2020) showed that both NCAM2 isoforms are expressed in the developing cortex and that its expression is necessary for correct neuronal migration, since NCAM2 knock-down leads to neuronal mispositioning. In the present analysis, we observed that at P0, most E15-born control neurons were present in the upper portion of the cortical plate and displayed a typical immature pyramidal neuron shape, with a main apical dendrite directed towards the marginal zone (**Fig. 6A-E**). In the case of E15-born NCAM2.2-overexpressing neurons, we observed an altered distribution with a significant reduction of neurons in the upper portion of the cortical plate (Bin 10) (**Fig. 6A-B**). E15-born NCAM2.1-overexpressing neurons also had a tendency to allocate below bin 10 (**Fig. 6A-B**). A synergistic effect was found when embryos were electroporated with both isoforms (NCAM2.1+NCAM2.2) simultaneously (**Fig. 6A-B**). Additionally, in contrast with NCAM2 depletion, NCAM2 overexpression apparently does not disrupt normal dendritic arborization at this stage.

**Figure 6.**
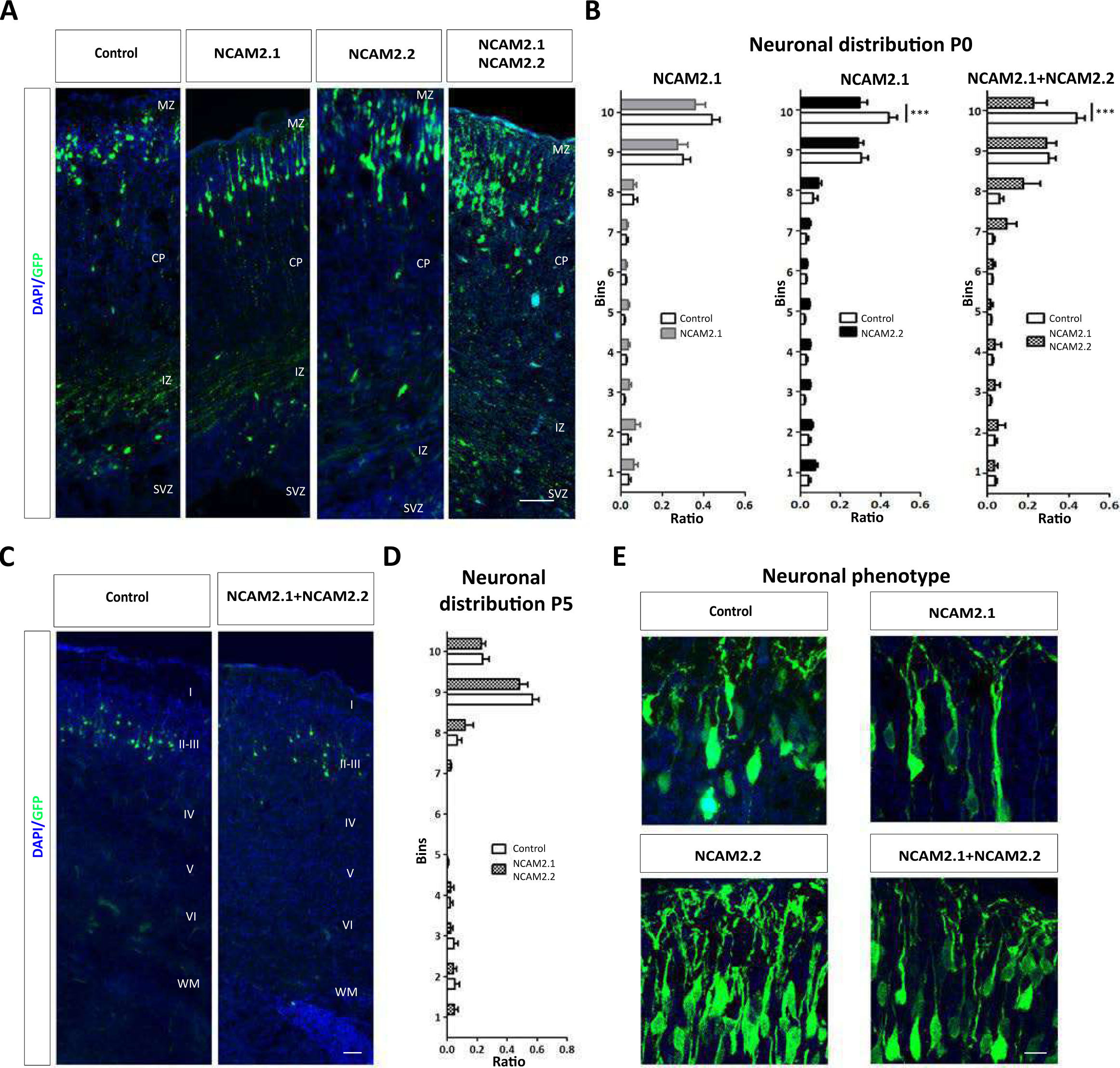
NCAM2 overexpression do not arrest embrionary RGPs but affects neuronal migration. **A)** Representative images from the reporter gene GFP in electroporated neurons in cortical sections from P0 mice. E15-born neurons were electroporated with control (left panel) and overexpression vectors. Sections were counterstained with DAPI. **B)** Distribution of transfected cells within cortical layers was quantified at P0 by dividing cortical thickness in 10 BINs. Data are presented as the ratio of neurons with somas located in each BIN. Overexpression of NCAM2.2 isoform or simultaneous expression of both isoforms (NCAM2.1+NCAM2.2) induce a reduced proportion of cells in the upper BIN. N=5-8 animals electroporated with control or overexpression constructs; *** P<0.001; two-way ANOVA, Bonferroni comparison *post hoc* test. **C)** Representative images from the reporter gene GFP in electroporated neurons in cortical sections from P5 mice. E15-born neurons were electroporated with control (left panel) and overexpression vectors for both isoforms (NCAM2.1+NCAM2.2; right panel). **D)** Distribution of transfected cells within cortical layers was quantified at P5 in 10 BINs. Data are presented as the ratio of neurons with somas located in each BIN. No differences were found within neuronal distribution between control and NCAM2-overexpressing neurons. N=6 electroporated animals with the constructs; two-ways ANOVA, Bonferroni comparison *post hoc* test. **E)** Higher magnification of representative images from transfected neurons at P0. Neurons show normal pyramidal neuronal morphology. CP, cortical plate; IZ, intermediate zone; MZ, marginal zone; SVZ, subventricular zone; I-VI, cortical layers. Scale bars: A,D) 50 μm; E) 10 μm.

In contrast, at P5, E15-electroporated neurons displayed a similar distribution in both for control and NCAM2-overexpressing conditions, with most neurons being located in the lower part of layer II-III (**Fig. 6C-D**). Our results suggest that NCAM2.1 and NCAM2.2 overexpression statistically not affect the proliferation, survival and differentiation of RGPs during embryonic stages but leads to transient migratory deficits.

### NCAM2 expression levels affect the growth of hippocampal-derived neurospheres

The implications of NCAM2 in adult neurogenesis were further investigated *in vitro* using neurospheres. Hippocampal NSCs were obtained from P6/7 mice and grown as neurospheres in medium containing EGF and bFGF. Neurospheres were dissociated and cells were infected with Control, ShNCAM2, NCAM2.1, or NCAM2.2-overexpressing viruses all of them co-expressing GFP as a reporter gene. GFP-positive cells were selected by flow cytometry, plated in 24 well plates and analyzed by ScanR microscopy to measure the individual area of a total of 100-300 growing neurospheres per condition during 5 consecutive days (**Fig. 7A**). Whereas the downregulation of NCAM2 led to the formation of larger neurospheres, compared to controls, neurospheres derived from NCAM2.1-or NCAM2.2-overexpressing cells tended to be smaller (**Fig. 7B-C**). Focusing the analysis on day 3, we observed a different distribution of the neurospheres according to their area. The descriptive analysis of the frequency distributions shows that the mean and median values of the distribution are lower in the NCAM2.1 and NCAM2.2 overexpressing neurospheres than in controls; and higher in the ShNCAM2 condition (**Fig. 7D-E**).

**Figure 7.**
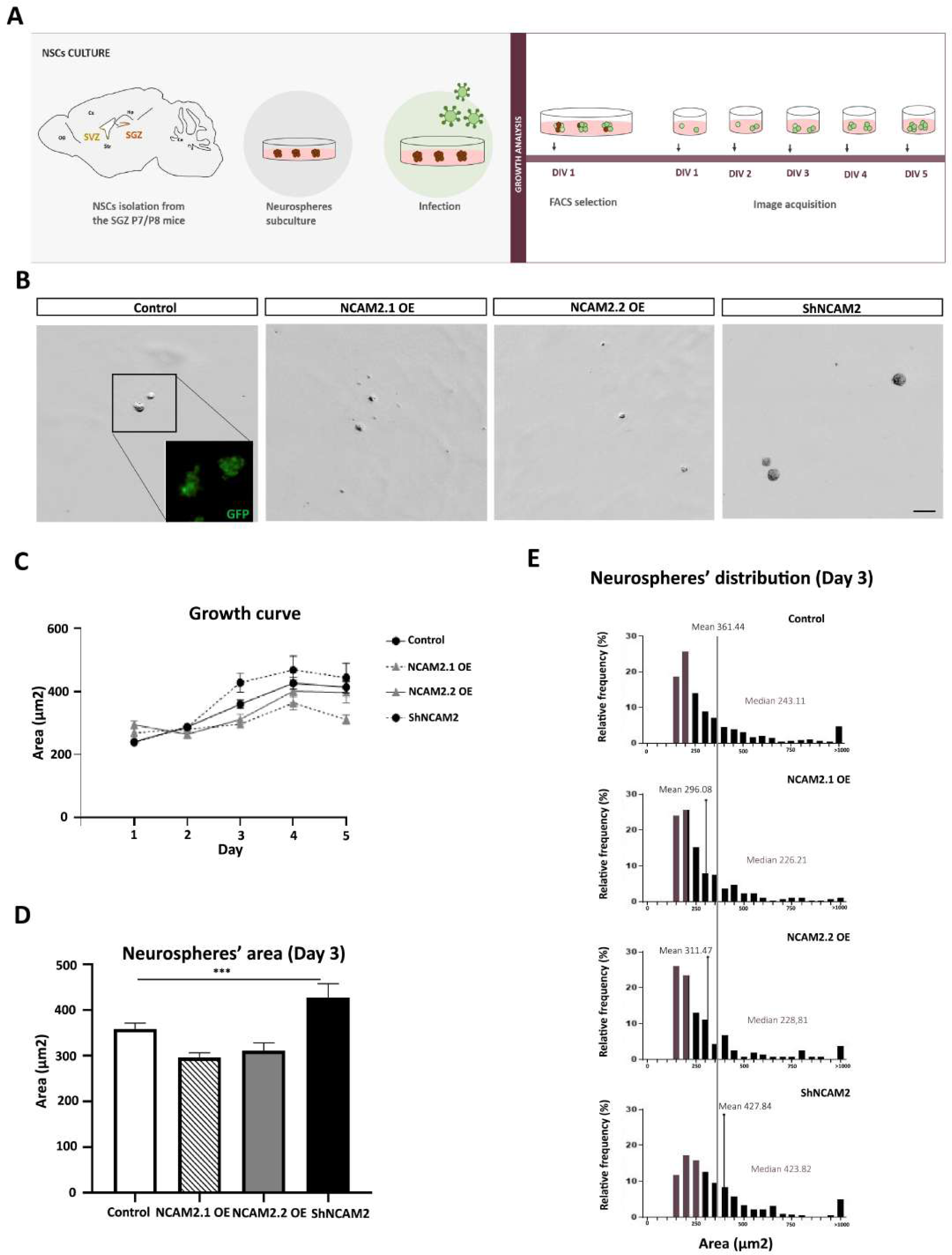
NCAM2 expression levels affect the proliferation of NSCs grown as neurospheres. A) Scheme showing the protocol for the obtention of post-natal mouse neurospheres from the neurogenic niches. Progenitor cells were isolated and grown as neurospheres A.1) Neurospheres where infected with control, NCASM2 overexpressing or ShNCAM2 viruses, selected by flow cytometry and plated in non-adherent plates. The area of the infected neurospheres was analysed by Scan-R microscopy for 5 consecutive days. **A.2)** Cells were plated in adherent coverslips, infected with control, NCAM2 overexpressing or ShNCAM2 viruses and maintained 5 days in differentiation conditions before fixation. B) Representative images of control, ShNCAM2, or NCAM2 overexpressing neurospheres after 3 days *in vitro*. **C)** Quantification of the time-course progress for the area of neurospheres for 5 consecutive days after sorting of infected cells. N= 100-300 neurospheres per condition, 1 independent experiment. **D)** Comparison of the area of neurospheres at 3 days *in vitro*. **E)** Histograms of control, NCAM2.1 OE, NCAM2.2 OE and ShNCAM2 neurospheres distribution according to their area 3 days after FACS selection. Coloured bars label percentile 50. Data are presented as mean ± SEM; Kruskal-Wallis test, *** P<0.001. Scale bar: B) 100 µm.

**Figure 8.**
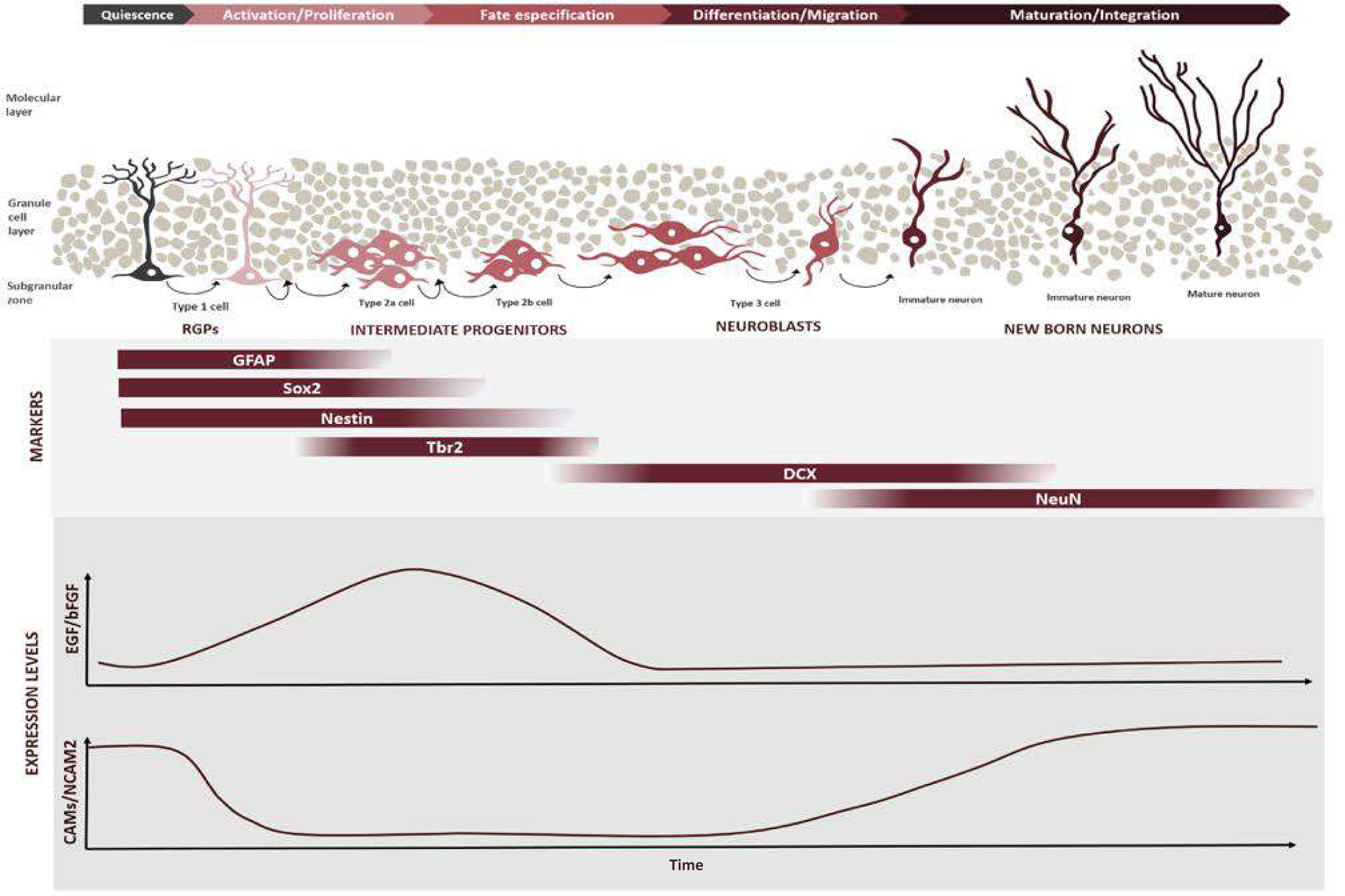
Model of RGPs regulation by NCAM2 expression levels in the hippocampus. Schematic representation of the proposed model for NSC regulation by NCAM2 expression. RGPs (Type I cells) are GFAP/Sox2/Nestin positive cells and are maintained in a quiescent state in the SGZ of the dentate gyrus. Upon activation, they generate Tbr2 positive proliferating intermediate progenitors (Type 2 cells). Those transit-amplifying progenitors produce neuroblasts (Type 3 cells) that express DCX and differentiate into NeuN positive granule cells. New-born neurons mature and become functional neurons of the hippocampal circuits. This process is regulated by different intrinsic and extrinsic factors, such as growth factors. We postulate that the levels of cell adhesion molecules such as NCAM2 protein are crucial for the regulation of NSC quiescence, the activation of proliferation and for the proper neuronal differentiation and maturation in later stages (Shin et al. 2015; Morizur et al. 2018; Parcerisas et al. 2020).

These findings further support the notion that NCAM2.1 and NCAM2.2 are involved in the regulation of NSCs proliferation.

## DISCUSSION

The present work provides a deeper understanding on the relevant functions of NCAM2 during embryonic development and adult neurogenesis. Our results suggests that NCAM2 levels regulate the RGP-to-immature neuron transition in the adult DG. In contrast, our data indicate that correct NCAM2 levels are not necessary for cortical neurogenesis, but relevant for cortical migration.

The injection of lentivirus to modulate the expression of NCAM2 in the progenitor cells of the SGZ in the hippocampus reveals a compelling role of NCAM2 in the regulation of neural progenitors. While the depletion of NCAM2 had minor effects, the overexpression of NCAM2 seems to arrest cells into an RGP-like phenotype and delay the formation of new granule cells, as characterized by morphology, immunohistochemical markers, and ultrastructure. However, when analyzing the effects of NCAM2 overexpression in the regulation of embryonic RGPs, we did not find clear evidences of any alterations in the survival, proliferation or differentiation of progenitor cells. We found that NCAM2 upregulation results in an early and transiently altered neuron distribution, suggesting a delay in their migration during cortical development. Our previous results also showed that the downregulation of NCAM2 led to an alteration of cortical migration leading to mislocalization of layer II-III fated neurons and altered morphology (Parcerisas et al., 2020). Neuronal migration is a key process in corticogenesis, the disruption of which is associated to many diseases including autism and schizophrenia (Hussman et al., 2011; Petit et al., 2015; Scholz et al., 2016). The mechanism underlying the effects of NCAM2 are not known. The interaction of NCAM2 with microtubule-associated proteins, such as MAP1B, that also participate in the regulation of neuronal migration has also been described (González-Billault et al., 2005; Kawauchi & Hoshino, 2008; Parcerisas et al., 2020; Parcerisas, Ortega-gascó, et al., 2021).

Focusing on the functions of NCAM2 in neurogenesis, our data suggest different roles of NCAM2 during adult and embryonic stages. In spite of the embryonic origin of adult RGPs, adult and embryonic progenitors are subject to distinct regulation (Urbán and Guillemot 2014; Berg et al. 2018; Daniel Berg et al. 2019). While embryonic RGPs have a highly proliferative rate necessary for the rapid growth of neural tissues (Urbán et al., 2019; Urbán 2014); adult RGPs are mostly found in a quiescent state, a mitotic-dormant phase with a low rate of metabolic activity but with a high sensitivity to environment signals (Urbán et al., 2019). The quiescence of RGPs is actively maintained and the regulation of the transition from quiescence to activation is crucial to preserve a pool of RGPs throughout life. Adult RGPs are found in neurogenic niches, specialized microenvironments composed by different cellular types, ECM molecules, soluble factors and cell surface molecules (Bian, 2013). Neurogenic niches are crucial for the regulation of RGPs properties and to maintain the quiescence/activation balance (Llorens-Bobadilla and Martin-Villalba, 2017; Basak et al., 2018; Kalamakis et al., 2019) as they convey the different physiological stimuli (Fabel and Kempermann 2008; Wang et al. 2011; N and F 2014; Ding et al. 2020) that induce the activation of quiescent RGPs.

Cell adhesion molecules are key elements of the neurogenic niches. They are important for sustaining the architecture of the niche but also participate in signal transduction regulating stem cells, survival, proliferation, migration or differentiation. As a matter of fact, different cell adhesion molecules such as cadherin/protocadherins, VCAM1, L1CAM or NCAM1 have been identified playing a distinct role in the neurogenic niches (K. Angata et al., 2007; Bian, 2013; Boldrini et al., 2018; Bonfanti, 2006; Dihné et al., 2003; Karpowicz et al., 2009; Marthiens et al., 2010; Morante-Redolat & Porlan, 2019; Morizur et al., 2018; Shin et al., 2015). Specifically, it has been described that cell adhesion molecules could be important regulators of the quiescence/activation balance. The genetic profiles of RGPs showed an enriched expression of genes involved in cell-microenvironment interaction and cell-cell adhesion, and genes linked to cell membrane (Artegiani et al., 2017; Basak et al., 2018; Ding et al., 2020; Dulken et al., 2017; Hochgerner et al., 2018; Llorens-Bobadilla et al., 2015; Morizur et al., 2018; Shin et al., 2015). Upon activation, RGPs proliferate and progress to rapid amplifying intermediate progenitors or type II cells. A decrease in the expression of some cell adhesion molecules seems to be necessary for the activation of quiescent RGPs, their transition to intermediate progenitors and the proliferation of these progenitors (Morizur et al., 2018; Shin et al., 2015; Codega et al., 2014; Xie et al., 2020). A similar expression pattern has been presented in this study when immunodetecting NCAM2 in the SGZ populations. The proposed pattern of NCAM2 expression along dentate neurogenesis cell types, supported by single cell RNA (Shin et al., 2015), confirms high NCAM2 expression in type I progenitors and low levels in intermediate progenitors. In fact, the expression pattern of *Ncam2* gene during the early neurogenic events is similar to other genes related to the maintenance of stem cells quiescence (e.g: NPas3 or Aqp4) (Shin et al., 2015; Urbán et al., 2019) presenting high levels of expression in qNSCs that progressively decrease during their activation and transition to intermediate progenitors (Shin et al. 2015; Morizur et al. 2018) (**Supplementary Fig. 2, Fig. 9**). Once the precursor cell phase is completed, the levels of NCAM2 seem to experiment a progressive increase in the newborn DCX positive maturing neurons reaching high levels of expression in NeuN neurons. The increase of NCAM2 could be explained by the relevance of the protein for dendrite development, axon formation and synaptogenesis (Alenius & Bohm, 2003; Kulahin & Walmod, 2010; Winther et al., 2012, Parcerisas et al., 2020).

The levels of NCAM2 seems to be important for the regulation of RGPs behaviour. In fact, our data show how changes in NCAM2 levels modifies the normal course of the neurogenic events. The upregulation of NCAM2 dramatically decrease the generation of newborn neurons. Diverse underlying mechanisms could explain these findings. The upregulation of NCAM2 could affect the survival of the newborn cells, induce the de-differentiation of developing neurons or either alter the differentiation of the newborn neurons. However, considering the expression pattern of the protein and the relevance of cell adhesion molecules in the regulation of RGPs (Codega et al. 2014; Morizur et al. 2018; Xie et al. 2020), our main hypothesis is that NCAM2 is important for the regulation and maintenance of RGPs quiescence. Considering that the overexpression of NCAM2 induces the retention of progenitor cells into a RGP state, we should expect that the downregulation of the protein promote the activation of RGPs to increase proliferation. In contrast, after inducing NCAM2 depletion in the hippocampus of injected mice, we did not detect an increase in the number of newly produced neurons. The underlying cause for this inconsistency might rest on the limitations imposed by the lack of uniformity in the infection of cells, preventing quantitative analyses of the number of new neurons generated. In order to overcome these limitations, we further investigated the effect of NCAM2 *in vitro* using a neurosphere assay. We observed that the downregulation of NCAM2 expression in progenitor cells *in vitro* increases the growth of neurospheres while overexpression of NCAM2 isoforms decreases the area of the neurospheres. The effects of NCAM2 in the proliferation of NSCs *in vitro* has previously been observed in progenitor cells that form the spinal cord (Deleyrolle et al. 2015) and supports the data obtained in the present study.

Taking these results together, we postulate that the regulation of NCAM2 expression levels is necessary for the maintenance of RGPs quiescence and the activation of proliferation. High levels of NCAM2 arrest cells in a quiescent state while the downregulation of *ncam2* allows RGPs to exit quiescence and enter the cell cycle to proliferate and differentiate (**Fig. 9**). The temporary retention of cells in the progenitor stages would led to a delay in the neurogenic events postponing the generation and maturation of granule cells although other explanations may contribute (e.g. changes in cell survival or differentiation to other cell types). Further research is needed to understand the mechanisms by which NCAM2 regulates RGPs quiescence, cell proliferation, and differentiation in adulthood. One hypothesis is that NCAM2 could interact with growth factor receptors such as the epidermal growth factor receptor (EGFR) or the fibroblast growth factor receptor (FGFR). Growth factors are important regulators of the activation of quiescent RGPs; for example, active RGPs in the SVZ could be identified by the expression of EGFR (Aguirre et al., 2010; Urbán et al., 2019). It has been described that NCAM2 binds to FGFR and EGFR (Deleyrolle et al. 2015; Rasmussen et al. 2018), and the interaction of other cell adhesion molecules, such as L1CAM or NCAM1, with FGFR has also been reported (Kulahin et al. 2008; Francavilla et al. 2009). Moreover, it has been shown that the overexpression of NCAM1 reduces baseline levels of EGFR, enhancing the EGF-induced receptor down-regulation, and that the depletion of NCAM2 increases the levels of the ErbB2 growth factor receptor (Povlsen et al. 2008; Deleyrolle et al. 2015). Another possibility is that NCAM2 expression could cause cytoskeletal rearrangements, which are known to influence the neurogenetic process (Compagnucci et al., 2016; Parcerisas et al., 2020; Parcerisas, Ortega-gascó, et al., 2021).

Neurogenic niches are complex microenvironments where RGPs receive and interact with multiple signals. Cell adhesion molecules are key elements for the transduction of the signals and the regulation of stem cells behavior. Our work provides evidence for a significant function of NCAM2 in the regulation of RGPs during adult neurogenesis. Furthermore, we reveal the importance of NCAM2 expression in the regulation of neuronal migration and differentiation during the corticogenesis process in the embryonic development. Overall, the present study contribute to a better understanding of the implications of NCAM2 during neuronal development and adult plasticity.

## CONFLICT OF INTEREST

The authors declare no competing financial interests.

## AUTHORS CONTRIBUTION

E.S., L.P., and A.P. conceived and designed the study. A.O-G. and A.P. performed most of the experiments and analyzed data. K.H. and S.S. designed and performed *in utero* electroporation. V.H-P. and J.M.G-V. designed and produced the electron microscopy experiments and analysis. F.U. supervised the RGP characterization. A.E-T and M.B participate in some experiments. A.O-G., A.P., V.H.-P., L.P., and E.S. wrote the manuscript. All authors read and corrected the manuscript.

## FUNDING

This work was supported by grants from the Spanish Ministry of Science, Innovation and Universities to V.H-P. (PCI2018-093062) and to E.S and L.P. (PID2019-106764RB-C21/ AEI/ 10.13039/501100011033), from the Spanish Ministry of health (ISCIII-CIBERNED), from The Secretary of Universities and Research of the Department of Economy and Knowledge of the Generalitat de Catalunya to A.P, from the Valencian Council for Innovation, Universities, Science and Digital Society (PROMETEO/2019/075) to J.M.G-V, and from the National Institute of Health (NIH) (R01NS109176) to S.S..

## ACKNOWLEDGEMENTS

We thank A. Lladó and S. Tosi (microscopy facility of the IRB-Barcelona) for FIJI macro design and technical assistance in ScanR acquisition; L. Bardia (microscopy facility of the IRB-Barcelona) for support and technical assistance; E. Coll and M. Calvo for technical assistance (microscopy facilities of the University of Barcelona). Marta Pérez for technical support in viral production (Universitat Internacional de Catalunya). The members of the Department of Cell Biology, Physiology and Immunology (University of Barcelona), specially J. Correas for cryostat support and Esther Verdaguer for valuable support; and members of the Soriano lab for experimental help and comments.

## ABBREVIATIONS

CAM: Cell adhesion molecules
CNS: Central nervous system
DAB: Diaminobenzidine
DAPI: 2-(4-amidinophenyl)-1H -indole-6-carboxamidine DG Dentate gyrus
EGF: Epidermal growth factor
EGFR: Epidermal growth factor receptor
FGF: Fibroblast growth factor
FGFR: Fibroblast growth factor receptor
GFAP: Glial Fibrillary acidic protein
GFP: Green Fluorescent Protein
GL: Granule layer
GPI: Glycosylphosphatidylinositol
H: Hilus
HRP: Horseradish peroxidase
IZ: Intermediate zone
L1CAM: L1 cell adhesion molecule
MAP2: Microtubule-associated protein 2
ML: Molecular layer
MTT: (3-(4,5-dimethylthiazol-2-yl)-2,5-diphenyltetrazolium bromide
NCAM1: Neural cell adhesion molecule 1
NCAM2: Neural cell adhesion molecule 2
NGS: Normal goat serum
NHS: Normal horse serum
NSC: Neural stem cell
PB: Phosphate buffer
PBS: Phosphate buffer saline
PFA: Paraformaldehyde
RGP: Radial glial progenitor
SGZ: Subgranular zone
Sox2: Sry-related HMG box transcription factor
SVZ: Subventricular zone
VCAM1: Vascular cell adhesion molecule 1

## Notes

### Competing Interest Statement

The authors have declared no competing interest.

