## Supplemental Information for "Regulation of adult neurogenesis and neuronal differentiation by Neural Cell Adhesion Molecule 2 (NCAM2)"

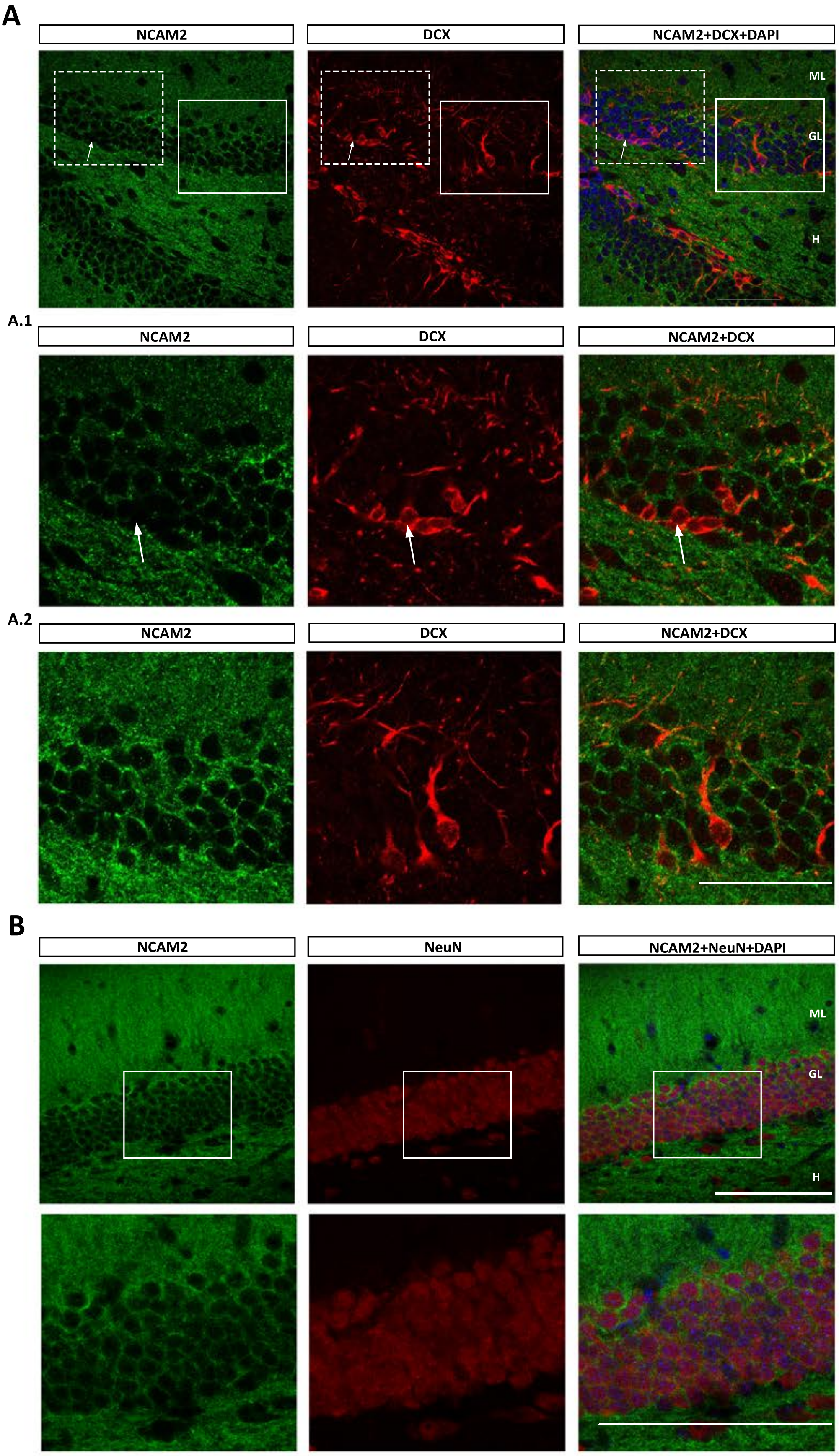


Supplementary Figure 1

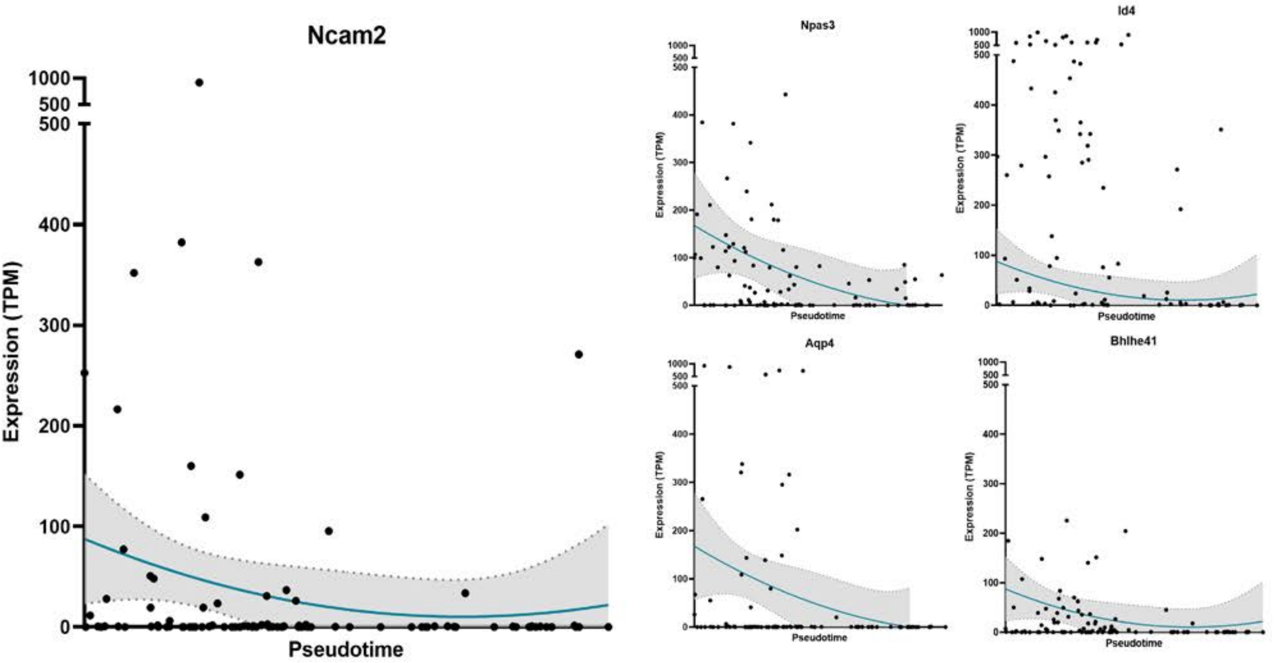


**Supplementary Figure 2**

### *Supplementary Information*

#### SUPPLEMENTARY MATERIALS AND METHODS

##### Representation of single cell genetic profiles

The genetic profiles of *Ncam2* and representative transcriptions factors were obtained from the supplementary data published in *Shin et al., 2015* (Table S6. Single-cell gene expression table according to the pseudotime progression). By single cell RNA seq, *Shin et al., 2015* revealed the expression of different genes during the "Pseudotime" progression. In the mentioned article, "Pseudotime" refers to the continuous progress from a quiescent neural stem cell (qNSCs) during their activation and transition to intermediate progenitors. The values corresponding to certain genes as appearing in *Shin et al., 2015* (Table 6) of single-cell gene expression during "Pseudotime" progression were adjusted to second order polynomial standard curves.

#### SUPPLEMENTARY FIGURES

##### Supplementary figure 1. Expression pattern of NCAM2 in the hippocampus.

**A)** Immunohistochemical characterization of NCAM2 expression in DCX positive cells in P45 mice hippocampus. A.1) Magnification of DCX positive/NCAM2 negative cells. A.2) Magnification of DCX positive/NCAM2 positive cells. Arrowheads label NCAM2/DCX positive cells**.** Arrows label NCAM2 negative/DCX positive cells**. B)** NCAM2 expression in NeuN positive cells in the hippocampus of P45 mice. ML: molecular layer; GL: granule layer; H: hilus. Scale bar: A, B) 20 µm; high magnifications 10 µm.

##### Supplementary Figure 2. Expression levels of *Ncam2* and representative transcription factors.

Gene expression levels represented as transcripts per million (TPM).along "Pseudotime" progression of *Ncam2* (left graph) compared to gene expression levels of representative transcription factors (Npas3, Id4, Aqp4, Bhehl41, right graphs) that are down-regulated during the activation of qNSCs and their transition to intermediate progenitors. Data obtained from *Shin et al., 2015* (Table S6. Single-cell gene expression table according to the pseudotime progression) was represented and fitted to second order polynomial standard curves with 95% confidence interval (gray area).
